# Velocity-selective arterial spin labelling bolus duration measurements: Implications for consensus recommendations

**DOI:** 10.1101/2024.05.04.592549

**Authors:** Ian D Driver, Hannah L Chandler, Eleonora Patitucci, Emma L Morgan, Kevin Murphy, Stefano Zappala, Richard G Wise, Michael Germuska

**Affiliations:** Cardiff University Brain Research Imaging Centre (CUBRIC), School of Psychology, Cardiff University, Cardiff, United Kingdom; Cardiff University Brain Research Imaging Centre (CUBRIC), School of Physics and Astronomy, Cardiff University, Cardiff, United Kingdom; Department of Neurosciences, Imaging and Clinical Sciences, ‘G. d’Annunzio University’ of Chieti-Pescara, Chieti, Italy; Institute for Advanced Biomedical Technologies (ITAB), ‘G. d’Annunzio University’ of Chieti-Pescara, Chieti, Italy

**Keywords:** Magnetic resonance imaging (MRI), perfusion, cerebral blood flow (CBF), arterial transit time (ATT), Fourier-transform velocity-selective inversion (FT-VSI), cerebrovascular reactivity (CVR)

## Abstract

Velocity-selective arterial spin labelling (VSASL) MRI is insensitive to arterial transit time. This is an advantage over other perfusion measurements, where long arterial transit times can lead to bias. Therefore, VSASL can be used to study perfusion in the presence of long arterial transit times, such as in the ageing brain, in vascular pathologies, and cancer, or where arterial transit time changes, such as during measurement of cerebrovascular reactivity (CVR). However, when calculating perfusion (cerebral blood flow, CBF, in the brain) from VSASL signal, it is assumed that images are acquired before the trailing edge of the labelled blood has arrived in the imaging slice. The arrival of the trailing edge of the labelled bolus of blood will cause an underestimation of perfusion. Here we measure bolus duration in young, healthy human brains, both at rest and during elevated CBF. Grey matter bolus duration was 1.61 ± 0.31 s, but there was a large spatial heterogeneity, with bolus duration being lower in anterior brain regions, with some areas having bolus duration < 1.2 s. We place these results in context of recommendations from a recent consensus paper, which recommends imaging 1.4 s after the label, potentially underestimating CBF in anterior regions. Further, we observed a 0.23 ± 0.12 s reduction in grey matter bolus duration with 5% CO_2_ inhalation. These results can be used to inform the experimental design of future VSASL studies, to avoid underestimating perfusion by imaging after the arrival of the trailing edge of the labelled bolus.

## 1. Introduction

Arterial spin labelling (ASL) MRI provides a non-invasive measurement of perfusion. It is being used to further our understanding of physiological mechanisms in both the brain (Haller et al., 2016; Liang et al., 2013; Mattsson et al., 2014; Satterthwaite et al., 2014; Segerdahl et al., 2015; Staffaroni et al., 2019) and body (Taso et al., 2023). Very generally, ASL consists of labelling water protons in arterial blood and waiting for the labelled water to perfuse into the tissue of interest before imaging. The original and predominant ASL method labels water protons based on spatial position, outside of the imaging volume (spatially-selective ASL) (Alsop et al., 2015; Dai et al., 2008; Detre et al., 1992; Edelman et al., 1994; Kim, 1995; Williams et al., 1992; Wong et al., 1998). This approach is dependent on waiting long enough for the labelled blood to flow from the labelled arteries into the tissue of interest, a time defined as the arterial transit time (ATT).

Recently, significant technical development has been made in another ASL method, termed velocity-selective ASL (VSASL), whereby water protons are labelled based on their velocity, rather than spatial position (Wong et al., 2006). The main advantage of this approach is that blood is labelled much closer to the tissue of interest, significantly reducing or eliminating ATT (Qin et al., 2022). This insensitivity to ATT makes VSASL a good candidate for studying mechanisms where ATT is long, or may change, which would result in a biased measurement of perfusion for spatially-selective ASL. Long arterial transit times have been observed in the ageing brain (Mahroo et al., 2024; Scheel et al., 2000), in vascular pathologies (Bokkers et al., 2009; Bolar et al., 2019; Fan et al., 2017; Qiu et al., 2012; Wang et al., 2013) and in gliomas (Qu et al., 2022). Changing ATT is also a source of bias in cerebrovascular reactivity (CVR) measurements made with spatially-selective ASL (Zhao et al., 2021). Therefore, there is a large range of applications that could benefit from the ATT insensitivity of VSASL.

VSASL consists of a velocity-selective labelling module followed by a vascular crushing module. The velocity-selective labelling module labels spins moving above a cut-off velocity (v_cutoff_). After a period where the labelled arterial blood spins perfuse into tissue, a vascular crushing module suppresses signal from spins moving above v_cutoff_, leaving the signal from perfused spins which decelerated as they perfused into the tissue. Here we define the time between the velocity-selective label and the vascular crushing module to be the Label-to-Crusher Time (LCT). The labelling module can either be formed of a train of saturation or inversion pulses. Labelling efficiency is higher with the inversion method and approaches that of pCASL (Qin et al., 2022). A Fourier transform velocity selective inversion (FT-VSI) labelling method (Qin & van Zijl, 2016) was chosen for this study, with FT-VSI recently demonstrating good test-retest reliability for measuring CBF in healthy participants (Xu et al., 2023). A recent consensus paper recommended LCT = 1.4 s and v_cutoff_ = 2 cm/s (Qin et al., 2022), so these were chosen as the starting point here.

The bolus duration is the time from the labelling module until the point when the trailing edge of the labelled bolus arrives in the tissue of interest. As no further fresh labelled blood arrives, the kinetic curve decays according to T_1_. Models used to quantify perfusion cannot account for an unknown timepoint where only fresh unlabelled blood is arriving, so the quantification will result in an underestimation in perfusion. Therefore, knowledge of bolus duration will improve specificity of perfusion measurements and will help to inform choice of LCT based on the bolus duration of the population to be studied. Bolus duration has been measured previously (Guo & Wong, 2015) as 2.03±0.08 s using velocity selective saturation ASL in five young healthy volunteers. However, that study did not focus on the spatial characteristics of bolus duration, whilst the early LCT space was sparsely mapped (values of [0.6, 1.2, 1.8, 2.4, 3.0, 3.6, 4.2] s used). A recent study raises questions over bolus duration (Xu et al., 2022), with FT-VSI ASL-based measurements of cerebrovascular reactivity (CVR) being underestimated by 40% when a LCT of 1.52 s was used, but when a shorter LCT of 0.5 s was used CVR was much closer to a phase-contrast based CVR measurement. This suggests that the bolus duration is shorter than 1.52 s, where CBF (especially during hypercapnia, where flow velocity increases) and CVR were underestimated. Previous studies generally consider bolus duration to be equivalent to LCT (the time between labelling and vascular crushing modules), such that the trailing edge of the labelled bolus is cut-off by the vascular crushing module. In this context, LCT and bolus duration are used interchangeably and are often referred to as *τ*. This assumes that the trailing edge does not reach the tissue of interest before the vascular crushing module. Therefore, we aim to measure bolus duration in young healthy participants and place these measurements in context of previous studies and recommended values of LCT.

## 2. Methods

### 2.1. Hardware and general acquisition parameters

Data were acquired on a Siemens 3T Prisma system with body transmit and 32 channel head receive. An MPRAGE T_1_-weighted anatomical scan was acquired for image registration and tissue segmentation (TR/TE/TI = 2100/3.19/850 ms; flip angle 8°; 1.1x1.1x1.0 mm, sagittal orientation (FHxAPxRL); phase partial Fourier 7/8).

A Fourier transform velocity selective inversion (FT-VSI) ASL sequence (Qin & van Zijl, 2016) was implemented with regional pre-sat (2 s pre-delay), composite refocusing pulses, velocity-compensated control, and a sym-BIR-8-based vascular crushing module. v_cutoff_ was defined as per (Qin et al., 2022) and values of 2, 3 and 4 cm/s were used in *Experiment 1*, whilst 3 cm/s was used in *Experiment 2*. The FT-VSI pulse train was 62 ms in length with a 7.8 ms gap between each 20-degree hard pulse, the rise time for the gradients in the FT-VSI pulse train was 0.35 ms. The gradient amplitude was altered linearly to achieve the desired v_cutoff_, with an amplitude of 18.75 mT/m used to achieve a 2 cm/s v_cutoff_ in the inferior-superior direction.

Three background suppression pulses were timed to ensure negative perfusion contrast at each LCT. The timings of the background suppression pulses were interpolated from the results presented in (D. Liu et al., 2020), inversion times are given in *Table 1*.

**Table 1:** Background suppression timings, delay from the end of the labelling module to the centre of the inversion pulse. LCT is the time from the labelling module to the vascular crushing module.

| LCT (ms) | Inversion 1 (ms) | Inversion 2 (ms) | Inversion 3 (ms) |
| --- | --- | --- | --- |
| 550 | 288 | 323 | 353 |
| 660 | 329 | 426 | 476 |
| 800 | 380 | 557 | 632 |
| 960 | 439 | 706 | 808 |
| 1150 | 508 | 833 | 1016 |
| 1390 | 596 | 1107 | 1275 |
| 1670 | 699 | 1368 | 1574 |
| 2000 | 820 | 1677 | 1919 |
| 2400 | 967 | 2050 | 2329 |
| 2900 | 1150 | 2517 | 2828 |
| 3500 | 1370 | 3077 | 3407 |

A 2D EPI readout was used, with 3.3x3.3mm in-plane resolution (64x64 matrix), 15 slices (6mm thickness, 3mm slice gap), TE = 12 ms and 6/8 partial Fourier. TR was set as short as possible, ranging from 3.5-6.3 s from the shortest to longest LCT. A 50 ms delay was applied between the vascular crushing module and the readout to reduce eddy currents and the post-label delay increased by 42.5 ms for each ascending slice. Four tag/control pairs were acquired for each LCT with an M_0_ acquired at the start of each set. The M_0_ included the sym-BIR-8-based vascular crushing module to cancel out T_2_ weighting in the perfusion quantification. LCT values will be specified in each respective experimental section below.

### 2.2. Experiment 1: Bolus duration measurements

Thirteen young, healthy participants (8 female, 5 male; aged 18-52, median 22, mean±stdev 28±11 years) gave written informed consent. The School of Psychology, Cardiff University Ethics Committee approved this study.

To calculate bolus duration, eleven LCT values of [0.55, 0.66, 0.8, 0.96, 1.15, 1.39, 1.67, 2, 2.4, 2.9, 3.5] s were acquired. This was repeated for three v_cutoff_ values of 2/3/4 cm/s, with the v_cutoff_ order balanced across participants (as there were thirteen participants, 5 participants had an order 2/3/4, whereas 4 participants each underwent orders of 3/4/2 and 4/2/3 cm/s).

### 2.3. Experiment 2: Bolus duration response to hypercapnia CVR

Nineteen young healthy participants (10 female, 9 male; aged 18-45, median 22, mean±stdev 26±9 years) gave written informed consent. The School of Psychology, Cardiff University Ethics Committee approved this study. The sample size for experiment 2 was powered a-priori based on the mean grey matter bolus duration measurements from experiment 1 and anticipating a reduction in bolus duration of 0.1 s with hypercapnia (based on a similar reduction in bolus arrival time measured in pCASL and PASL previously (Donahue et al., 2014; Ho et al., 2011; Su et al., 2017). For an effect size of 0.7, alpha = 0.05 and power = 0.8, a total sample size of 19 was required (G*Power 3.1).

The gas delivery system used was introduced previously by our group (Germuska et al., 2016), with mass flow controllers (MKS Instruments, Wilmington, MA, USA) delivering either 25 L/min of medical air (normocapnia) or 5% CO2 (hypercapnia). Participants wore a close-fitting face mask (Quadralite, Intersurgical, Wokingham, Berkshire, UK) attached to a breathing circuit following the design of Tancredi and colleagues (Tancredi et al., 2014). Exhaled gas was monitored throughout, with a sampling line from the face mask to a rapidly responding gas analyzer (PowerLab, ADInstruments, Sydney, Australia).

To calculate bolus duration, a set of eight LCT values of [0.55, 0.66, 0.8, 0.96, 1.15, 1.39, 1.67, 2] s were acquired. This set was repeated four times, [normocapnia – hypercapnia – normocapnia – hypercapnia]. The last three LCT values (2.4, 2.9 & 3.5 s) acquired in *Experiment 1* were omitted here, to reduce the amount of time a participant was at hypercapnia in each set. Each set only started once end-tidal PCO_2_ stabilised to a steady state (∼ 1 minute following the transition). A 3 cm/s v_cutoff_ was chosen to reduce cerebrospinal fluid (CSF) signal contamination, which is a problem for hypercapnia, where global vasodilation displaces CSF volume (Blockley et al., 2011; Thomas et al., 2013).

### 2.4. Data analysis

FT-VSI ASL data were concatenated across LCT values for each dataset, then motion correction was performed (MCFLIFT, FSL (Jenkinson et al., 2002)). Bolus duration and CBF were estimated using a standard 1-compartment kinetic model and unconstrained minimisation (Quasi-Newton method).

The ASL signal was modelled as per (Qin et al., 2022)

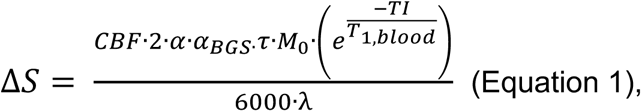

Where ΔS is the signal difference between the label and control images. *τ* = LCT for the case where LCT <= bolus duration. However, *τ* = bolus duration in the case where LCT > bolus duration (i.e. the image is acquired after arrival of the trailing edge of the labelled bolus). TI is the time between labelling pulse and readout (i.e. LCT + slice timing delay relative to the vascular crushing module for each slice), CBF is the cerebral blood flow (ml/100g/min), α is the inversion efficiency (0.56), α_BGS_ is the additional signal attenuation caused by the three background suppression pulses (0.86), λ is the blood-tissue-partition coefficient (0.9), M_0_ is the signal intensity of the proton density image, and T_1,blood_ = 1.6 s is the T1 value of blood (Lu et al., 2004).

M0 scans were coregistered to the MPRAGE using FSL epi_reg (Greve & Fischl, 2009; Jenkinson et al., 2002) and the MPRAGE coregistered to MNI152 space (Fonov et al., 2011) using FSL FLIRT (Jenkinson et al., 2002).

Grey matter maps were segmented using FSL BET (Smith, 2002) and FAST (Zhang et al., 2001) on the MPRAGE. Grey matter partial volume estimate maps were realigned into native ASL space. All grey matter analyses were performed using a mask of voxels with a grey matter partial volume estimate of at least 50%. An additional threshold was applied to the M0 of greater than 50% of the maximum signal to avoid areas where the EPI signal had significant signal drop-off (intra-voxel dephasing close to air spaces).

The Harvard-Oxford Cortical Atlas (Desikan et al., 2006), which is provided in FSL in MNI152 space, was realigned into native ASL space. Regional analyses were performed within the grey matter mask defined in the previous paragraph by defining each region based on a 25% threshold of that region’s partial volume estimate (Craddock et al., 2012).

Group average bolus duration and CBF maps were calculated in MNI152 space by realigning parameter maps into MNI152 space at 2mm isotropic resolution, then taking the mean across participants.

## 3. Results

### 3.1. Experiment 1a: bolus duration measurements

Results are first presented in the context of how bolus duration can inform choice of sequence timings (LCT, the labelling to vascular crushing delay time), presenting data from all three v_cutoff_ values in parallel. Secondly, results are presented comparing v_cutoff_ values, in context of the impact of v_cutoff_ on the data and how these results can inform the choice of v_cutoff_ for CVR experiments, where CSF signal contamination can bias results.

Grey matter average bolus duration (mean ± standard deviation across participants) was 1.61 ± 0.31 s / 1.55 ± 0.28 s / 1.50 ± 0.28 (2/3/4 cm/s v_cutoff_). All participants’ grey matter bolus duration, with median and interquartile range are shown in *Figure 1a*. Grey matter average CBF results are shown in *Figure 1b*. Note that there is a strong negative correlation (r = -0.8; p = 10^-3^) across participants between CBF and bolus duration, consistently across the three v_cutoff_ values, with participants with high CBF having shorter bolus duration than those with lower CBF.

**Figure 1:**
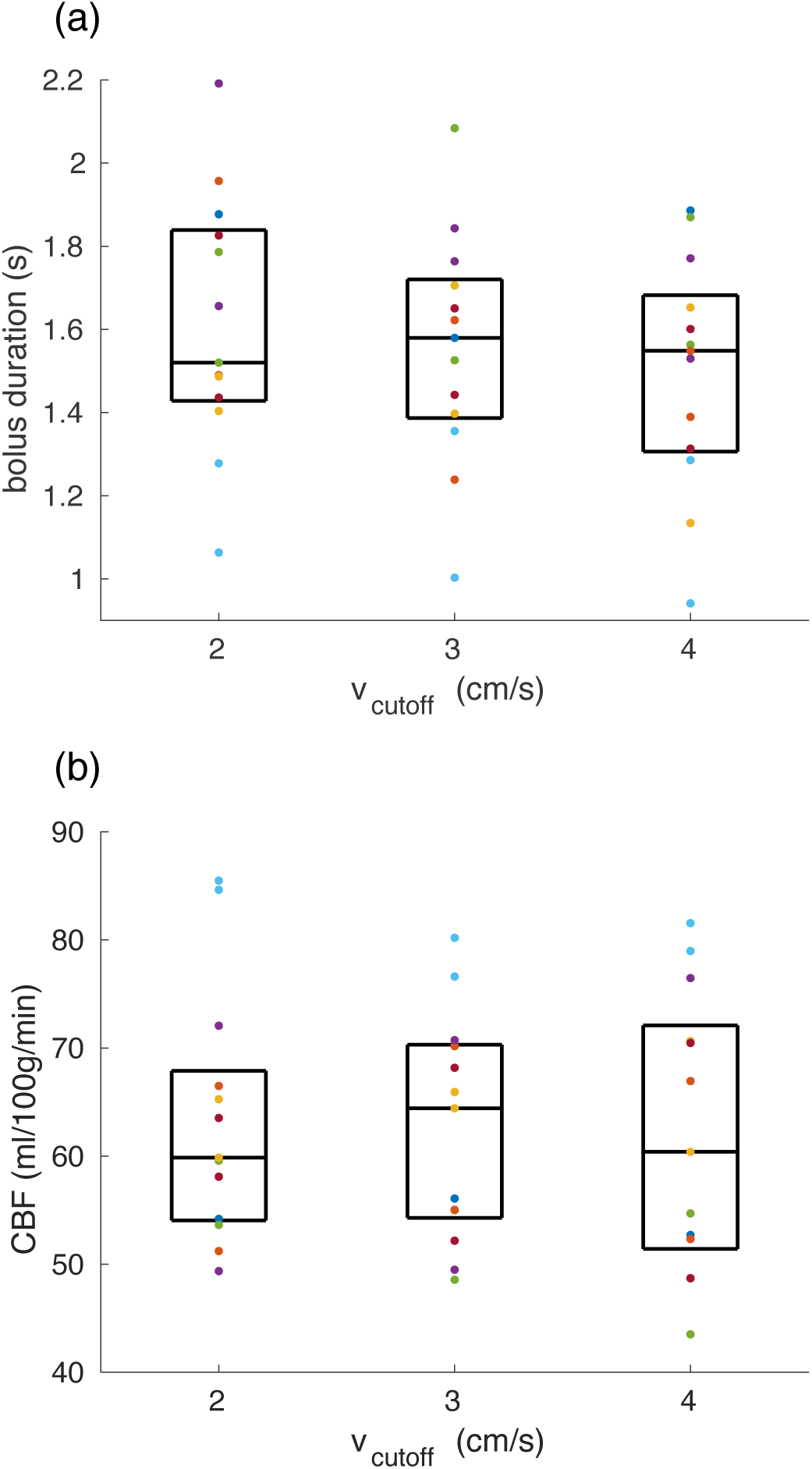
(a) Grey matter mean bolus duration for each vcutoff (box shows median and interquartile range, points show each participant’s data). (b) Grey matter mean CBF

Bolus duration maps in *Figure 2* show a large amount of spatial heterogeneity, with anterior regions having shorter bolus duration than posterior regions. This is further investigated in *Figure 3*, showing grey matter average bolus duration sub-divided into regions defined by the Harvard-Oxford Cortical Atlas (Desikan et al., 2006), with a representative dashed line at 1.4 s showing that many regions have bolus duration shorter than the consensus recommended LCT = 1.4 s (Qin et al., 2022).

**Figure 2:**
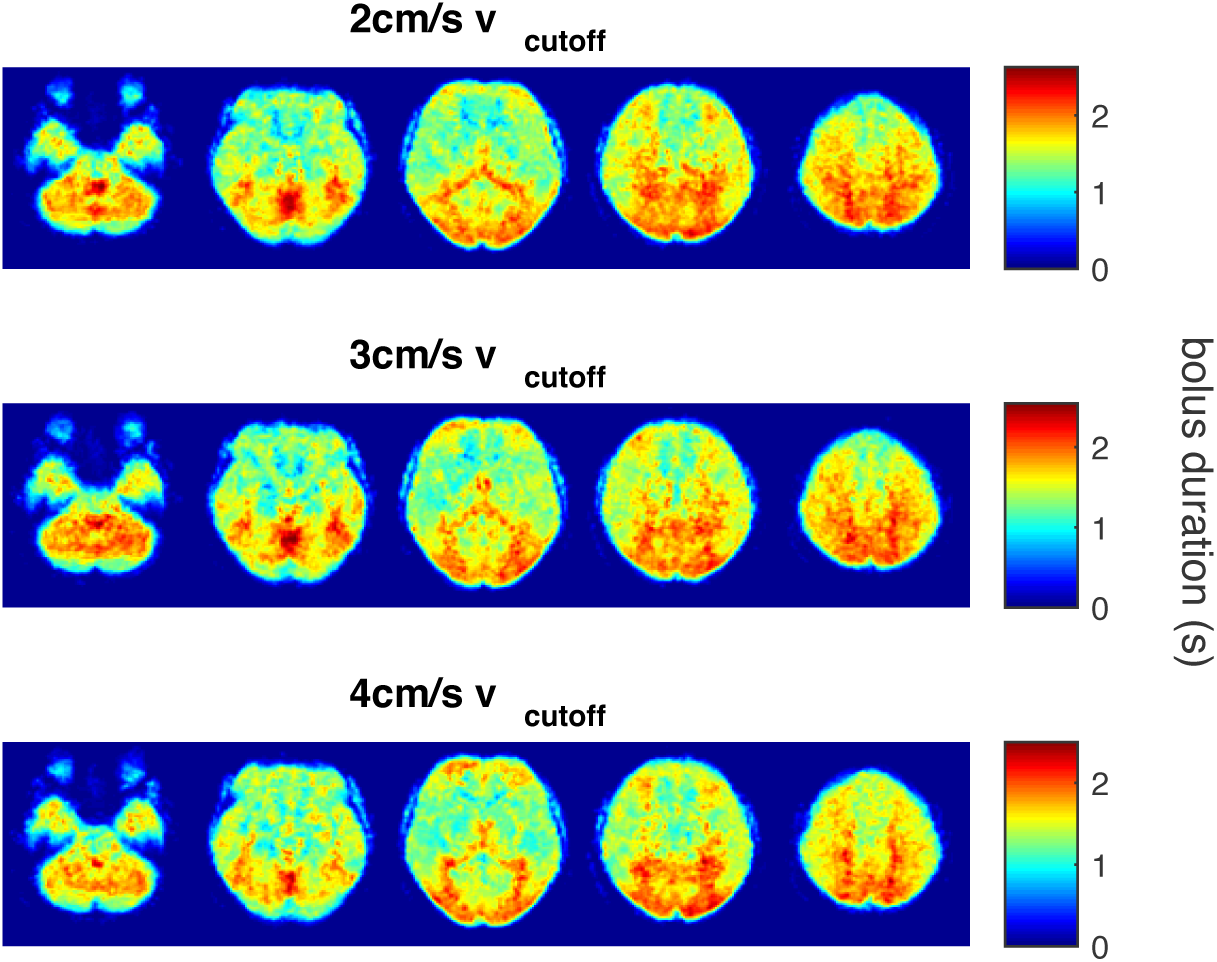
Bolus duration maps, averaged across all participants and shown for each vcutoff. The five slices shown are at MNI152 z = -32, -12, 8, 28, 48 mm.

**Figure 3:**
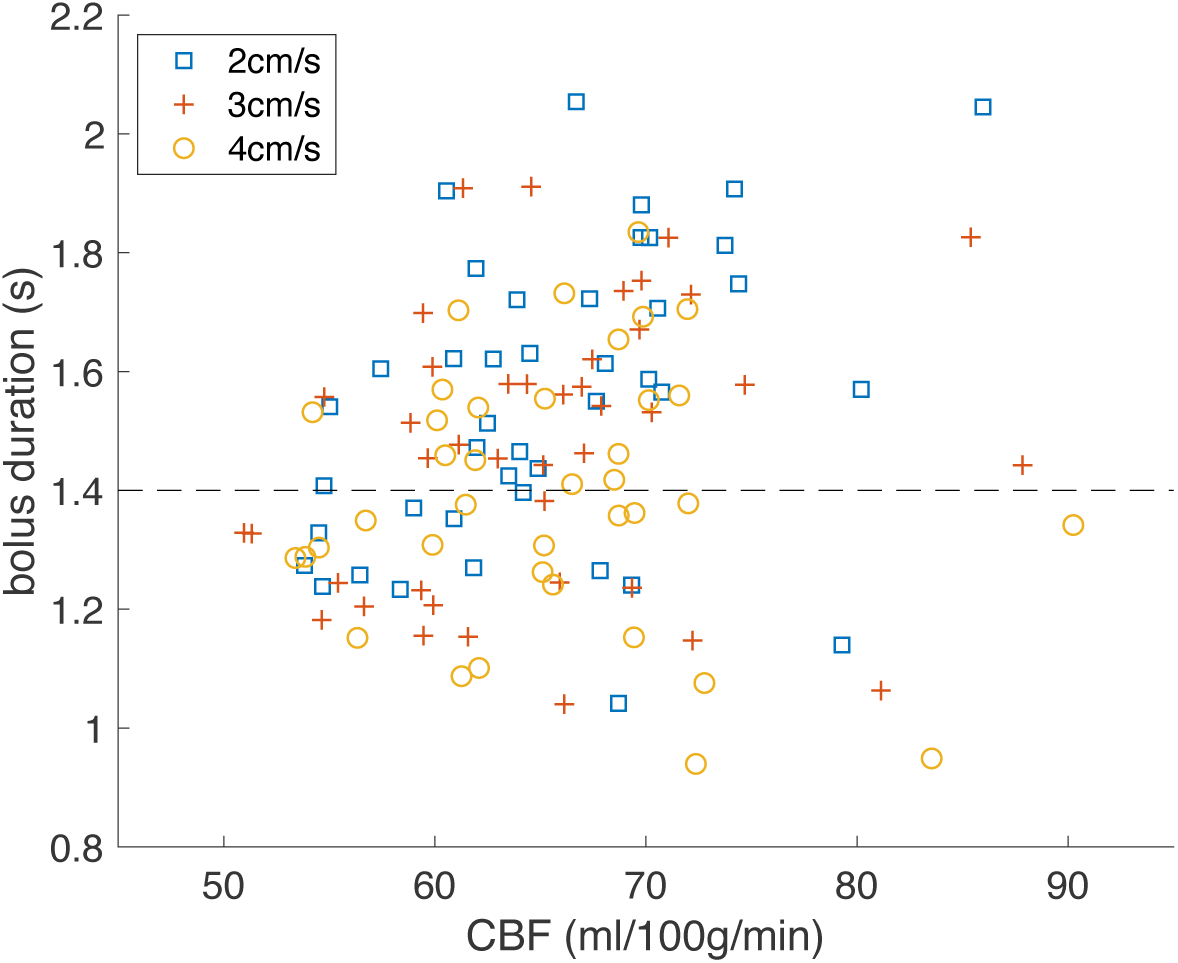
Scatter plot comparing bolus duration with CBF. Each point shows a data from a region in the Harvard-Oxford Cortical Atlas, averaged over all participants. Data for each vcutoff is shown by a different marker shape and the dashed horizontal line shows a bolus duration of 1.4 s.

To inform the choice of LCT in future experimental design, the percentage of Harvard-Oxford Cortical Atlas regions (out of 48) for which bolus duration is below a range of values of LCT are reported in *Table 2*. Dropping from a LCT of 1.4 s to 1 s brings the fraction of regions with bolus duration less than LCT (so at risk of underestimating CBF) from approximately 40% to less than 10 %.

**Table 2:** Percentage of the 48 Harvard-Oxford Cortical Atlas regions which have a bolus duration less than the labelling to vascular crushing delay time (LCT) for a range of LCT times and for each vcutoff value. Data is presented as median [interquartile range] % across participants.

| LCT (s) | 2 cm/s $v_{\text{cutoff}}$ | 3 cm/s $v_{\text{cutoff}}$ | 4 cm/s $v_{\text{cutoff}}$ |
| --- | --- | --- | --- |
| 1.4 | 38 [17-55] % | 38 [24-63] % | 44 [24-71] % |
| 1.2 | 21 [10-27] % | 19 [6-34] % | 21 [10-48] % |
| 1 | 2 [2-8] % | 8 [4-17] % | 6 [2-19] % |
| 0.8 | 0 [0-2] % | 0 [0-3] % | 0 [0-5] % |

### 3.2. Experiment 1b: impact of v_cutoff_ for designing CVR experiments

Reviewing the above results in the context of minimising signal contamination from CSF spaces, *Figure 4* demonstrates that the apparent perfusion signal from CSF spaces is reduced from 2 cm/s to 3 cm/s v_cutoff_ and reduced to a lesser extent from 3 cm/s to 4 cm/s v_cutoff_. *Figures 1-3* suggest that if there are any differences in CBF and bolus duration measured at each v_cutoff_, then they are small compared to intra- and inter-participant variation. Therefore, for CVR experiments, where CSF spaces are apparently displaced by vasodilation, a higher v_cutoff_ than the consensus recommendation 2 cm/s (Qin et al., 2022) is more appropriate.

**Figure 4:**
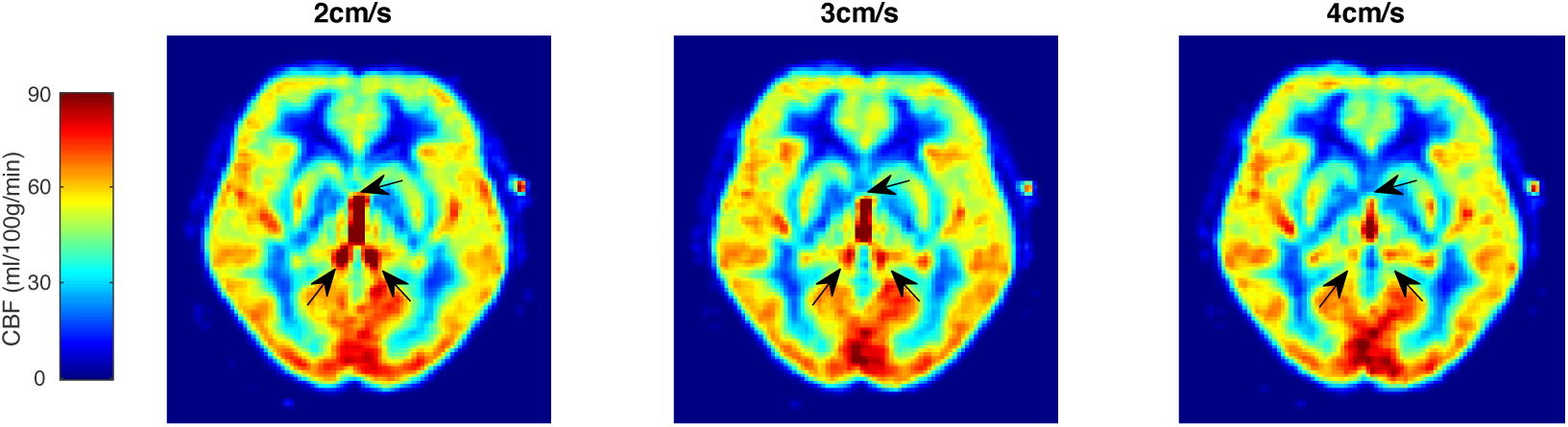
A single slice (MNI152 z = +2 mm) of the group-average CBF map, highlighting the effect of vcutoff on CSF signal contamination. Black arrows show CSF signal from the lateral ventricles at the same position for each vcutoff.

### 3.3. Experiment 2: Bolus duration response to hypercapnia CVR

The hypercapnic challenge caused an 8.3 ± 2.4 mmHg increase in P_ET_CO_2_. This caused mean grey matter bolus duration to decrease by 0.23 ± 0.12 s (paired 2-tailed t-test p = 9x10^-8^; t(18) = 8.5) and CBF to increase by 66 ± 33 % (p = 8x10^-8^; t(18) = 8.6). Absolute values for bolus duration and CBF are shown for each participant, with median and interquartile range in *Figure 5*. Note that the values for bolus duration are lower than for *Experiment 1*. This is due to an upper limit on absolute bolus duration estimation of the longest LCT value (2 s for Experiment 2) leading to longer values being clipped to 2 s in this experiment. This does not affect the relative changes measured. Group average maps of baseline bolus duration and change in bolus duration with 5% CO_2_ are shown in *Figure 6*. The reduction in bolus duration appears to be restricted to grey matter.

**Figure 5:**
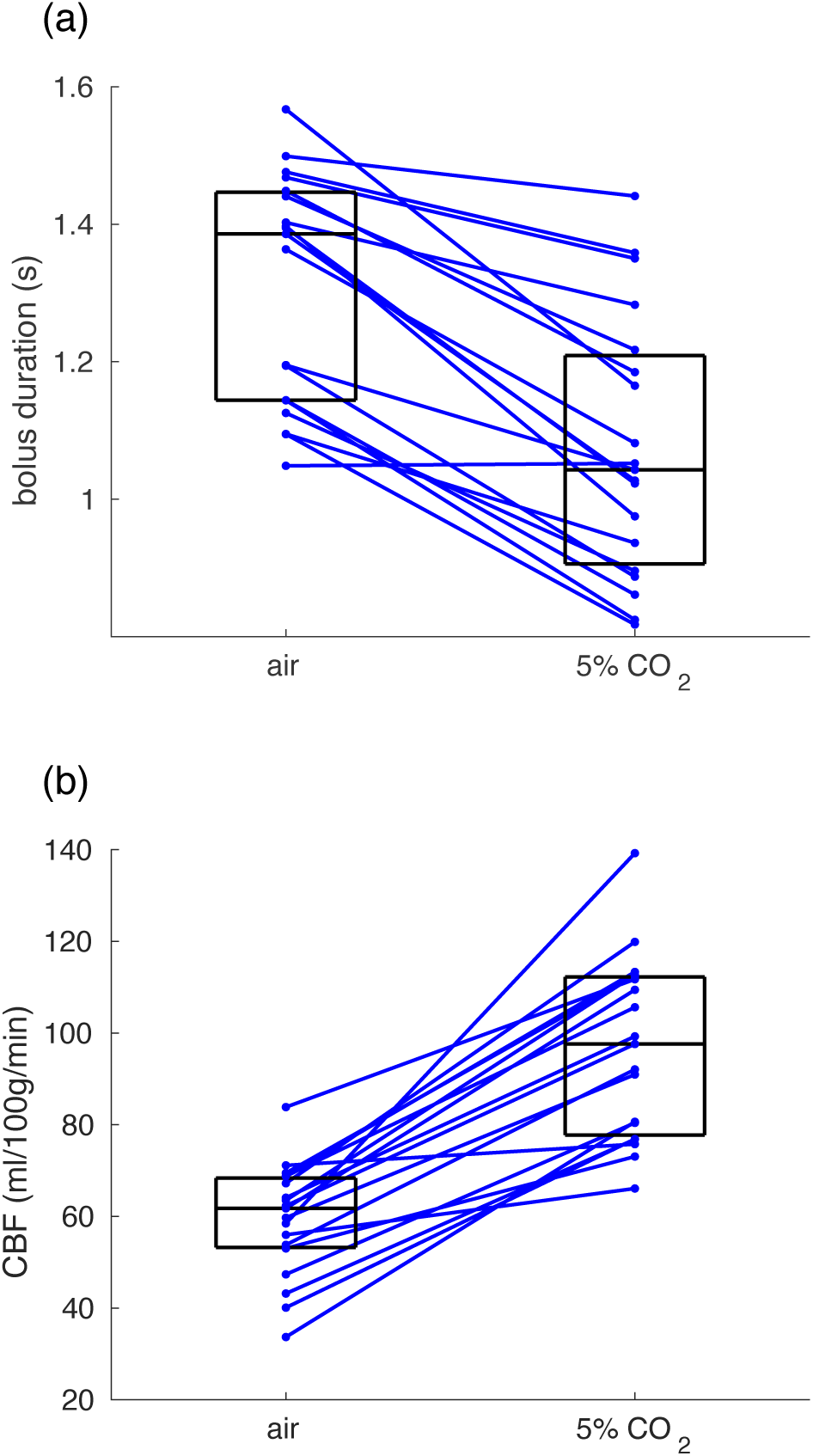
The change in grey matter mean (a) bolus duration and (b) CBF with 5% CO2. Lines match participants’ values in air and 5% CO2 breathing conditions and boxes show median and interquartile range across participants.

**Figure 6:**
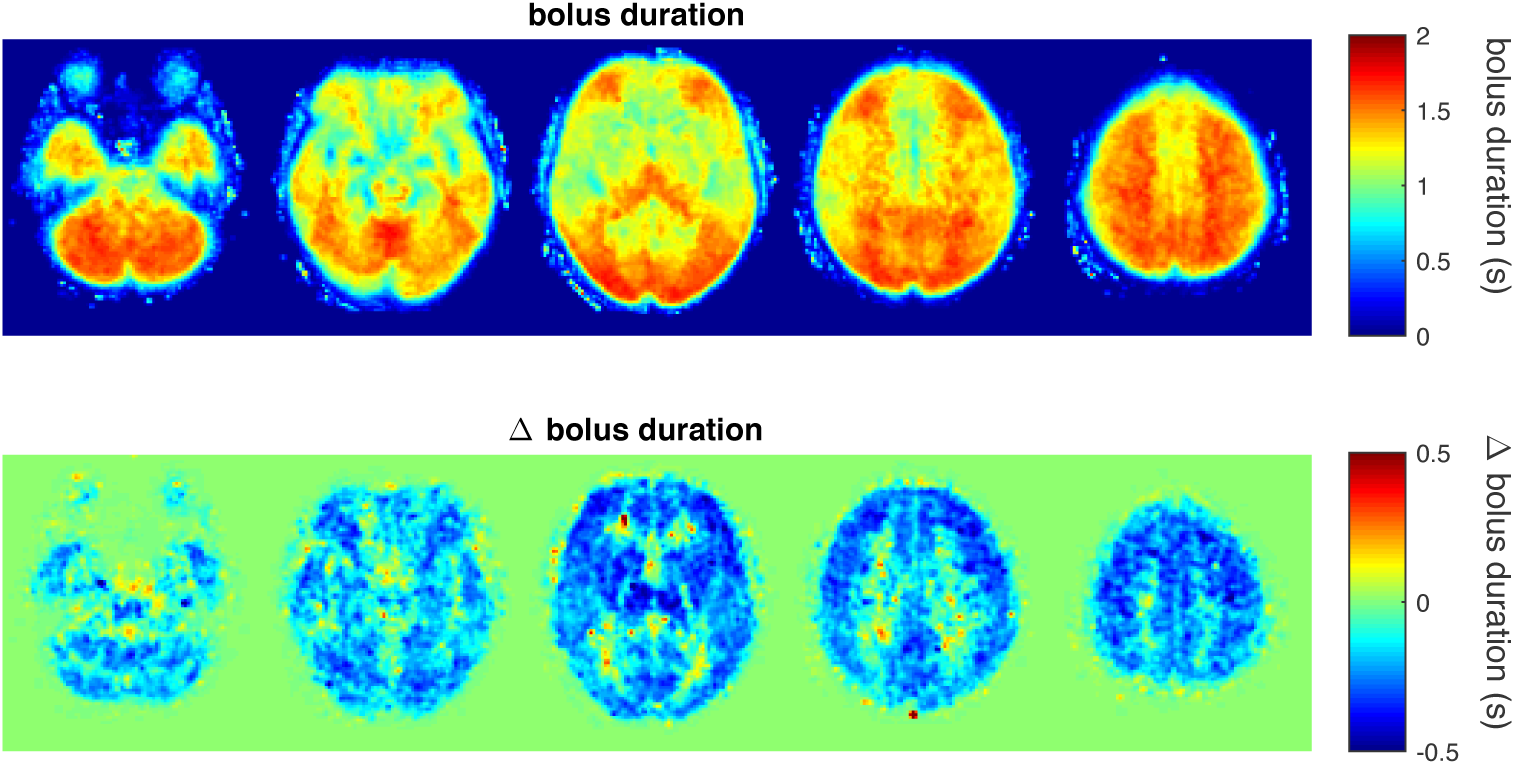
Top row – bolus duration maps during air breathing (averaged across all participants). Note the reduced colour scale compared to Figure 2. Bottom row – group average maps of the change in bolus duration with 5% CO2 hypercapnia. The five slices shown are at MNI152 z = -32, -12, 8, 28, 48 mm.

## 4. Discussion

This work presents measurements of bolus duration in the young healthy brain for Fourier-transform velocity selective inversion ASL. We show that whilst grey matter average bolus duration is 1.6 s, there is a large spatial variability, with anterior regions having shorter bolus durations than posterior regions. We extend these findings by measuring a reduction in bolus duration of 0.2 s during hypercapnia in an experiment designed to measure CVR, providing an example where a physiological challenge can change bolus duration.

These results start to put an upper bound on the time between label and vascular crusher (LCT), such that LCT should be shorter than the bolus duration to avoid underestimating CBF by missing the trailing edge of the labelled bolus. The motivation for longer LCT is that more labelled blood can perfuse into tissue, increasing signal. Therefore, the optimal LCT is a balance of specificity (shorter LCT) and sensitivity (longer LCT). Future VSASL studies can take these results to inform the choice of LCT based on bolus duration for their population and region of interest. For example, in the young, healthy, predominantly female cohort studied in *Experiment 1*, a choice of LCT of 1.5 s would be appropriate for studying posterior brain regions, but to obtain an unbiased measurement of CBF in anterior brain regions, an LCT of 1 s would be required. The shorter bolus duration in anterior regions is consistent with shorter ATT in the anterior circulation, compared to the posterior circulation, that has been observed using pCASL (Pinto et al., 2023; Zhao et al., 2021). To put a scale on the potential to underestimate CBF through using LCT longer than the bolus duration, Xu and colleagues recently reported a 40% underestimation in CBF CVR measured using FT-VSI ASL with a 1.52 s LCT (Xu et al., 2022).

The arrival of the trailing edge of the labelled bolus determines bolus duration. The trailing edge is caused by B0 and B1 inhomogeneities distal to the area of focus (below the neck for the brain). Velocity encoding of fast flowing spins is less effective in the presence of field inhomogeneities, cropping the labelled bolus. The extent to which the trailing edge of the labelled bolus is cropped (and hence bolus duration) depends on the choice of both labelling and control methods. Bolus duration might be extended by use of sinc-VSI pulses, which are less sensitive to B0 inhomogeneities than the rect-VSI pulses used here (Guo et al., 2021). Further, in this study we used a velocity-compensated control, which may be more affected by distal B0 and B1 inhomogeneities than a velocity-insensitive control. Therefore, bolus duration may be different for a velocity-insensitive control. The choice of velocity-selective saturation or inversion is also likely to affect bolus duration, as saturation of fast-moving spins will be less sensitive to field inhomogeneities than inversion. For example, grey matter average bolus duration was measured previously as 2.03±0.08 s with velocity-selective saturation (Guo & Wong, 2015). The longer bolus duration could be due to the saturation method, compared to the inversion method used in our study. Other factors that could contribute to the differences between the two studies are the sparser sampling of LCT, or if there were a gender difference in bolus duration, with the previous study having mostly male participants, whereas *Experiment 1* here has mostly female participants. Unfortunately, the sample size is too small to search for gender differences in bolus duration. Our current study was designed to measure bolus duration with a FT-VSI method matching recent consensus recommendations (Qin et al., 2022), so the bolus duration measurements will be widely applicable to future studies using the consensus implementation.

Secondary to measuring bolus duration, *Experiment 1* also investigated the effect of v_cutoff_ on the FT-VSI ASL signal. Minimizing v_cutoff_ brings the label into smaller arteries and arterioles, closer to the capillary bed. This extends the lead edge of the labelled bolus and minimises the contribution of small blood vessels to the subtracted ASL signal. However, signal contamination from CSF is present with low v_cutoff_ values. Whilst there appears to be a small, non-significant reduction in bolus duration with v_cutoff_ changing from 2 to 4 cm/s (*Figures 1-3*), this is small compared to intra- and inter-participant variability in bolus duration, whilst CBF measurement was not affected by v_cutoff_. However, increasing v_cutoff_ above 2 cm/s reduced CSF signal contamination (*Figure 4*). Therefore, we chose to use a v_cutoff_ of 3 cm/s for *Experiment 2*, where CSF contamination could confound measurement of CBF CVR, where CSF spaces are displaced by global vasodilation (Blockley et al., 2011; Thomas et al., 2013). In the case where a patient group may have atrophy, or for CVR studies (Zhao et al., 2021), our results support using a v_cutoff_ of 3 cm/s to reduce CSF contamination, whilst not significantly reducing the size of the labelled bolus.

Further, in *Experiment 2* we measured a 0.2 s reduction in bolus duration in response to 5% CO_2_. This is consistent with increased blood flow velocity during hypercapnia and the 40% underestimation of CBF CVR reported by Xu and colleagues when using FT-VSI ASL with a 1.52 s LCT (Xu et al., 2022). The trailing edge of the labelled bolus arrives 0.2 s earlier during hypercapnia than normocapnia. For regions with a bolus duration less than 1.52 s, this reduction will cause an additional underestimation of CBF over that at normocapnia, whilst other regions, with normocapnic bolus duration close to 1.52 s, CBF will be underestimated for hypercapnia, but not normocapnia. In both cases, CBF CVR will be underestimated when using a 1.52 s LCT. In our CVR experiment, we measure a CBF CVR of 8 %/mmHg. This is approximately double that measured using pCASL (P. Liu et al., 2019), but is at the upper bound of phase-contrast whole brain CBF CVR measurements (Taneja et al., 2020; Xu et al., 2022), which are used to validate ASL.

A limitation of our method for measuring bolus duration is that we assume a sharp trailing edge of the labelled bolus which does not account for any dispersion. Dispersion would smooth out the kinetic curve. The effect of dispersion in the data, not accounted for in the model kinetic curve, would be to overestimate bolus duration.

In conclusion, heterogeneity in bolus duration in FT-VSI can lead to underestimation of CBF with the currently recommended LCT of 1.4 s. Our data suggests that a LCT of 1s or less will minimise any bias in CBF measurements, although at the expense of reduced signal-to-noise in superior and posterior brain regions. The use of a shorter LCT will be particularly important when studying groups or states where bolus duration is expected to change, such as cerebrovascular reactivity mapping. We also found that increasing v_cutoff_ from 2 to 3 cm/s did not have a significant effect on either bolus duration or CBF quantification. However, there is a clear reduction in CSF contamination, which if present could bias cerebrovascular reactivity measurements, or confound CBF quantification in certain pathologies. Therefore, we recommend increasing FT-VSI v_cutoff_ to at least 3cm/s for such studies.

## 5. Data and code availability

Participants only gave ethical approval for anonymised data to be shared, so we are unable make the raw ASL or MPRAGE data publicly available. Instead, we provide bolus duration and CBF parameter maps for each participant, in group (MNI152) space, and summary grey matter and Harvard-Oxford atlas region-averaged data, available at 10.17605/OSF.IO/6SJNQ. The analysis code for estimating CBF and bolus duration is available online 10.5281/zenodo.11065798.

## 6. Author Contributions

**Ian Driver:** Conceptualization, Methodology, Formal analysis, Investigation, Writing - Original Draft. **Hannah Chandler:** Investigation, Writing - Review & Editing. **Eleonora Patitucci:** Investigation, Writing - Review & Editing. **Emma Morgan:** Investigation, Writing - Review & Editing. **Kevin Murphy:** Funding acquisition, Writing - Review & Editing. **Stefano Zappala:** Investigation, Writing - Review & Editing. **Richard Wise:** Methodology, Funding acquisition, Writing - Review & Editing. **Michael Germuska:** Conceptualization, Methodology, MRI sequence development, Formal analysis, Investigation, Funding acquisition, Writing - Review & Editing.

## 8. Acknowledgements

This research was funded by a Wellcome Trust Strategic Award [104943/Z/14/Z]. For the purpose of open access, the author has applied a CC BY public copyright licence to any Author Accepted Manuscript version arising from this submission.

IDD, HLC and KM are supported by the Wellcome Trust [WT224267]

MG is supported by the Wellcome Trust [220575/Z/20/Z] and the Engineering and Physical Sciences Research Council [EP/S025901/1]

RGW is supported by

European Union-NextGenerationEU-Italian Ministry of University and Research (MUR), National Plan for Recovery and Resilience (PNRR) and Projects of National Relevance (PRIN), Project Code: P2022ESHT4, Project Title: “Advancing MRI biomarkers of brain tissue microstructure and energetics in Multiple Sclerosis.” Funding call No. 1409 of 14.09.2022, Concession decree No. 1367 of 01.09.2023 adopted by MUR, ERC Panel LS5 “Neuroscience and Disorders of the Nervous System”. CUP: D53D23019210001

European Union – NextGenerationEU under the National Plan for Recovery and Resilience (PNRR), Mission 4 Component 2 – M4C2, Investment 1.5 – Call for tender No. 3277 of 30.12.2021 Italian Ministry of Universities Award Number: ECS00000041, Project Title: “VITALITY - Innovation, digitalization and sustainability for the diffused economy in Central Italy,” Concession Decree No. 1057 of 23.06.2022 adopted by the Italian Ministry of University and Research. CUP D73C22000840006.

